# Dental aging offers new insights to the first epigenetic clock for common dolphins (*Delphinus delphis*)

**DOI:** 10.1101/2025.07.20.665818

**Authors:** Eva-Maria F. Hanninger, Katharina J. Peters, Livia Gerber, Ashley Barratclough, Emma L. Betty, Emily I. Palmer, Steve Horvath, Karen A. Stockin

## Abstract

Determining exact age in wild odontocetes is essential for understanding population dynamics, survival, and reproduction, yet remains logistically challenging. While epigenetic aging is emerging as a valuable approach, only nine species-specific clocks currently exist. Most have been calibrated using captive known-age animals or well-studied wild populations. Only two previous studies have used dental ages from stranded or bycaught individuals. This is due to concerns that dental age inaccuracies, especially in older animals, may affect epigenetic clock performance. To explore this, we developed the first species-specific epigenetic clock for common dolphins (*Delphinus delphis*), analysing DNA methylation at 37,492 cytosine-phosphate-guanine sites in skin samples from stranded and bycaught dolphins with estimated dental ages. Elastic net models with Leave-One-Out Cross-Validation were applied to three subsets: the ‘*relaxed*’ subset (all individuals; *n* = 75, median absolute error (MAE) = 2.02, *r* = 0.81, R^2^ = 0.66), the ‘*strict*’ subset (excluding individuals with minimum dental age estimates only; *n* = 73, MAE = 2.29, *r* = 0.81, R^2^ = 0.66), and the ‘*restricted*’ subset (excluding outliers with prediction errors > 6 years; *n* = 63, MAE = 1.80, *r* = 0.91, R^2^ = 0.82) to compare performance. Our models consistently underestimated the age of dolphins >16 years, even when minimum dental ages were applied, suggesting absolute errors between dental and epigenetic estimates unlikely reflect dental aging error. Additionally, post-mortem decomposition condition code (DCC 1 to 3) did not affect age prediction, signalling promise for future epigenetic clocks calibrated with strandings and bycaught individuals.

## 1. Introduction

In wildlife management, age determination plays a vital role in understanding population structure, an individual’s reproductive state, and lifespan (Barratclough, Sanz-Requena, et al., 2019; Betty et al., 2022). Such knowledge of population age structure provides crucial insights into population viability and resilience to both anthropogenic and environmental pressures (Betty et al., 2019; Heydenrych et al., 2021; Manlik et al., 2022; Palmer et al., 2022). More broadly, understanding life-history features is essential for effective conservation and management of protected species (Moore & Read, 2008), particularly those impacted by human-induced threats such as fisheries interactions, habitat degradation, and pollution (Murphy et al., 2018; Peltier et al., 2021; Stockin et al., 2010; Tulloch et al., 2020). However, accurate age estimates are essential to ensure reliable assessment of key life-history parameters, such as age at sexual maturity and inter-calving intervals (Manlik et al., 2016; Palmer et al., 2022; Verborgh et al., 2021). Yet obtaining such estimates is particularly challenging in odontocetes (toothed whales and dolphins), which exhibit limited external signs of aging (Barratclough et al., 2023; Beal et al., 2019; Krzyszczyk & Mann, 2012). While total body length can assist in assessing age class (i.e., neonate/juvenile/adult), growth varies greatly among individuals of the same species and age, making it unreliable for precise age estimation (Betty et al., 2019; Chivers, 2018). Instead, age estimation requires either long-term field studies (Connor & Krützen, 2015; Peters et al., 2023) or invasive methods, such as tooth extraction (Lockyer, 1993; Westgate & Read, 2007).

Typically, odontocetes are aged by quantification of growth layer groups in teeth, also known as ‘tooth aging’ (e.g., Betty et al., 2022; Evans et al., 2002; Lockyer, 1993, 1995; Maas, 2009; Murphy et al., 2014; Westgate & Read, 2007). Tooth aging requires trained, experienced readers and calibration using individuals of known age (Perrin & Myrick, 1980). While growth layer groups are generally assumed to represent annual deposition in most species — with the exception of beluga whale *Delphinapterus leucas* (e.g., Goren et al., 1987; Waugh et al., 2018) — their interpretation becomes increasingly challenging in older individuals. Contributing factors include varying degrees of tooth wear, compression of growth layer groups, and the presence of accessory lines (additional growth lines that may not correspond to annual deposition; Barratclough et al., 2023; Murphy et al., 2014; Read et al., 2018). Additionally, due to its invasive nature, tooth extraction is typically restricted to post-mortem investigations, except for procedures conducted under anaesthesia (Barratclough, Wells, et al., 2019), and accordingly does not permit assessment of demography in free-ranging populations (Betty et al., 2023). Consequently, age estimates are often derived from stranded or bycaught individuals, which may not accurately represent age structure of free-ranging populations and may bias population viability analyses and survivorship estimates (Betty et al., 2023).

Emerging molecular methods offer a minimally invasive alternative that can be applied to free-ranging cetaceans, using samples such as skin or blood, for example, to assess epigenetic patterns of DNA methylation for age estimation (Bocklandt et al., 2011; Horvath, 2013a; Horvath & Raj, 2018; Teschendorff & Horvath, 2025). Due to a phenomenon called ‘epigenetic drift’, genomic regions gain or lose cytosine methylation with age (Issa, 2014; Poulsen et al., 2007; Sen et al., 2016).

The Mammalian Methylation Consortium carried out epigenome wide association studies of age in 348 mammalian species including mysticetes and odontocetes (Haghani et al., 2023; Lu et al., 2023). The correlation between chronological age and DNA methylation at CpG sites (cytosine-phosphate-guanine) enables the development of epigenetic clocks, which are regarded as the most reliable predictors of age available (De Paoli-Iseppi et al., 2017; Guevara & Lawler, 2018; Jylhävä et al., 2017; Simpson & Chandra, 2021). The initial development of epigenetic clocks requires a highly accurately known age calibration population (Mayne et al., 2023). To date, eleven epigenetic clocks exist for odontocetes: three for common bottlenose dolphin (*Tursiops truncatus*; Barratclough et al., 2021; Beal et al., 2019; Robeck, Fei, Haghani, et al., 2021), two for Indo-Pacific bottlenose dolphin (*T. aduncus*; Peters et al., 2023; Yagi et al., 2023), one for killer whale (*Orcinus orca*; Parsons et al., 2023), one for beluga whale (Bors et al., 2021), one for Hector’s and Māui dolphins (*Cephalorhynchus hectori hectori* and *C. h. maui*; Hernandez et al., 2023), one for Risso’s dolphin (*Grampus griseus*; Mori et al., 2024) and two multi-species odontocete or cetacean clocks, which were cross-validated with nine (Robeck, Fei, Lu, et al., 2021), respectively 13 species (Zoller et al., 2025).

Most odontocete clocks have been calibrated on animals in human care (Barratclough et al., 2021; Robeck, Fei, Haghani, et al., 2021; Robeck, Fei, Lu, et al., 2021), or long-term observational studies of well monitored populations (Beal et al., 2019; Parsons et al., 2023; Peters et al., 2023; Yagi et al., 2023). At the time of our study, only three prior studies used growth layer groups of stranded and bycaught animals for initial calibration of their epigenetic clocks (Bors et al., 2021; Hernandez et al., 2023; Mori et al., 2024). Notably, concerns have been raised about whether errors in dental aging may affect the robustness of epigenetic clocks, particularly in older individuals where growth layer groups can be confused with accessory lines (Bors et al., 2021; Hernandez et al., 2023; Mori et al., 2024; Zoller et al., 2025).

Here, we present the first epigenetic clock for common dolphins (*Delphius delphis*) — an essential tool for future population viability analysis (PVA) in a species highly impacted by human activities (Peltier et al., 2016; Piroddi et al., 2011; Stockin et al., 2009; Tulloch et al., 2020). Further, we provide three distinct models that enhance current understanding of the reliability of dental aging for age calibration of methylated clocks. Our clock was developed using dental age estimates from individuals spanning the full known lifespan of free-ranging common dolphins, estimated at approximately 30 years (Perrin, 2009), with reports of individuals in human care reaching >35 years (Murphy et al., 2014). As dental aging can be less accurate in older individuals due to tooth wear and growth layer group compression, this approach allows us to evaluate how such uncertainties can affect clock accuracy. Specifically, we hypothesised that: (1) a clock model excluding individuals with uncertain dental age estimates would outperform one that includes all samples, and (2) a model based on all samples would perform worse than previously published odontocete clocks calibrated using known-age individuals from long-term observational studies or human care.

## 2. Materials & Methods

### 2.1. Sample collection

As part of ongoing longitudinal studies on health and life-history, we accessed a 30-year tissue archive of aged common dolphins spanning 0 to 34 years of age, for which sex, body length, sexual maturity, reproductive stage and radiographic bone aging had been completed *a priori*. Dental age was assessed via growth layer groups in the dentine of thin, decalcified, and stained sections of teeth by at least two independent experienced readers over three reads. If no agreement was reached, an additional tooth was prepared and examined by at least two readers until consensus was reached on the age estimate. In total, each animal had 1 to 3 teeth processed for aging. Age estimates were further validated against additional indicators of physical and sexual maturity, including total body length, the number of ovarian *corpora albicans* scars based on Palmer et al., (2022) and pectoral flipper radiographic aging (adapted from Barratclough et al., 2019). These metrics allowed cross-verification of age estimates across multiple biological parameters.

To ensure an adequate sampling size for the construction of a reliable epigenetic clock (Mayne et al., 2021), we selected skin from 84 common dolphin samples examined between 2000 and 2023. Sample selection was guided by multiple criteria including dental age (Supplementary Material, Table S1) and sex ratio (females: *n* = 47; males: *n* = 37). We included individuals with dental ages spanning the maximum known lifespan of common dolphins, with age and sex distribution represented equally as the sample availability would allow. Our samples originated from various mortality contexts, including bycatch (*n* = 9) and strandings (single stranding events: *n* = 46; mass stranding events: *n* = 28) and one individual from human care. Only animals of decomposition condition category DCC 1-3 (Ijsseldijk et al 2019) were included in the study (Supplementary Material, Table S1 and S2).

For 79 common dolphins, exact dental ages were ascertained. For the remaining five samples, only age ranges could be estimated. Here, we applied the minimum year of each age estimate to reduce the risk of age overestimation due to accessory lines. The age range of our samples spanned the estimated lifespan of the common dolphin, with the oldest free-ranging individual having a minimum dental age of 31 years. Additionally, we included a skin sample from a dolphin in human care (*Kelly*, Napier Marineland, held 1974 to 2008). Due to a pulp anomaly, only an age range could be determined, but Kelly was known to have spent 34 years in human care (Murphy et al., 2014).

### 2.2. DNA extraction

Genomic DNA was extracted from skin samples using a Quick-DNA™ Miniprep Plus Kit (Zymo) following the protocol for solid tissue samples. DNA purity was assessed using a NanoDrop Microvolume Spectrophotometer (Thermo Fisher Scientific) by an automated measurement of absorbance at the wavelengths 260 and 280 nm (Desjardins & Conklin, 2010). DNA was considered pure with an absorbance ratio ∼1.8 at 260nm/280nm (Desjardins & Conklin, 2010). DNA was purified using a DNA Clean & Concentrator Kit (Zymo) following the manufacturer’s instructions and concentration was measured using a QUBIT 4 fluorometer (Thermo Fisher Scientific) aiming for a total DNA amount per sample of >250ng.

### 2.3. DNA methylation data

A custom Infinium methylation array (HorvathMammalMethylChip40) assembled with 37,492 CpG sites was used measure DNA methylation (Arneson et al., 2022). The array targets highly conserved CpG sites across mammals allowing the assessment of DNA methylation in each mammal (Arneson et al., 2022). Sample quality control followed the Clock Foundation pipeline (Zoller & Horvath, 2024). Methylated (M) and Unmethylated (U) channels reporting background signal levels were marked as failed with the p-value threshold of *p* > 0.05. Additionally, the proportion of failed probes was counted for each sample. Samples with more than 1% of failed probes were considered fraction outliers. Further quality control steps included principal component analysis (PCA) performed on the centred, unnormalized beta-value matrix to identify major sources of variance in the dataset (Mishra et al., 2017), as well as hierarchical clustering analysis, whereas samples with higher than 2 standard deviations from the mean were marked as outliers (Zoller & Horvath, 2024). Nine samples were excluded from further analysis (see Results and Supplementary Material, Table S2).

### 2.4. Subsets

We developed the epigenetic clock using three distinct sample subsets. Details of sample selection for each subset are provided in the Supplementary Material (Table S1), in addition to age distributions by sex (Figure S1).

The first model, referred to as the ‘*relaxed*’ subset (*n* = 75; age range: 0–34 years) included all samples, also incorporating two individuals for whom only a minimum estimated dental age was available, both originated from individuals >30 years of age. The second model, termed *‘strict’*, excluded these two individuals and only incorporated animals with confirmed dental ages (*n* = 73; age range: 0–31 years). Following initial analyses, a third model was generated using only samples from the ‘*relaxed*’ subset that showed an absolute error of less than 6 years between actual and predicted age. This *‘restricted’* subset (*n* = 63; age range: 0–26 years) automatically excluded the two individuals with minimum age estimates. Except for three samples, all excluded individuals were ≥16 years old.

### 2.5. Age prediction

#### i) Elastic net regression with LOOCV

To build the epigenetic clocks, we used the `MammalMethylClock` R package which incorporates functionalities for the assessment, development and utilization of epigenetic clocks (Zoller & Horvath, 2024).

The package uses pre-existing life-history data from the Animal Aging and Longevity Database (AnAge^1^). In the case of *Delphinus delphis*, the AnAge data (female average age at attainment of sexual maturity (ASM): 4.8 years, male ASM: 4.3 years) did not align with previously published life-history data for common dolphins (Danil & Chivers, 2007; Murphy & Rogan, 2006; Palmer et al., 2022, 2023; Westgate & Read, 2007), therefore we manually overwrote the data with an average value of 8.15, as reported for male and female reproduction in Aotearoa New Zealand common dolphins (female ASM: 7.5 years, male ASM: 8.8 years; Palmer et al., 2022, 2023). For the gestation period, we used 1.05 years which was determined based on Aotearoa New Zealand common dolphins (Palmer et al., 2022). Based on these life-history features, we applied a log-linear transformation to our methylation data (Lu et al., 2023; Zoller & Horvath, 2024). This transformation accounts for accelerated epigenetic aging processes before attainment of sexual maturity (ASM) providing a logarithmic function if age is lower than age at sexual maturity (ASM) and a linear form in case age is greater than ASM (Lu et al., 2023). We further used the function `fun_llin2.inv`, to take species-specific gestation times into account.

To build a species-specific epigenetic clock, the package uses an elastic net regression (Zoller & Horvath, 2024), which is an algorithm for fitting generalized linear models with elastic-net penalties (Friedman et al., 2010). The elastic net algorithm works on very large datasets in which the number of predictor variables exceeds the number of observations (Friedman et al., 2010). Elastic nets are particularly useful in situations where there are many correlated predictor variables. We used an elastic net regression model (`glmnet` package) to determine the optimal mixing parameter (α) via cross-validation on 20 random subsets (each with 2/3 of the data). Models were fit across α values from 0 to 1 (in 0.1 steps), and the α with the lowest mean squared error was selected (Barratclough et al., 2021). Based on this analysis, the elastic net was chosen (‘*relaxed*’ subset: α = 0.4, ‘*strict*’ subset: α = 0.5, ‘*restricted*’ subset: α = 0.7) to remove any degeneracies and wild behaviour caused by extreme correlations (Friedman et al., 2010; Lu et al., 2023; Zoller & Horvath, 2024).

Cross-Validation was completed with a Leave-One-Out Cross-Validation (LOOCV) analysis (Zoller & Horvath, 2024). This function uses n-fold internal cross-validation, with *n* being the number of samples (Zhang, 1993; Zou & Hastie, 2005). For each sample of the data, the LOOCV approach skips one sample, fits the epigenetic clock on the remaining data and predicts the age of the omitted sample (Zhang, 1993; Zou & Hastie, 2005).

#### ii) Hybrid epigenetic clock

For the ‘*relaxed*’ subset, we applied a hybrid DNA methylation age prediction approach integrating a random forest classifier (RFC) and two elastic net regression (ENR) models following Barratclough et al., (2024). Physical maturity status was classified using an RFC with 1000 trees, trained to distinguish physically mature dolphins (≥18 years for females, ≥20 years for males, Palmer, 2023). Two ENR models were trained to predict log-linearly transformed ages, anchored on sex-specific sexual maturity thresholds (7.5 years for females, 8.8 years for males; Palmer et al., 2022, 2023). The first and second ENR was trained on all individuals and physically mature dolphins, respectively, to refine age predictions in older animals. For both ENRs, the optimal balance between LASSO and Ridge penalties (α) and the penalty parameter (λ) were selected by nested 5-fold internal cross-validation on the training data. Overall model performance was evaluated using 5-fold cross-validation. In each fold, the RFC and ENR models were retrained on the training data, and age predictions were generated as a weighted sum of the two ENR models, where the weight for the mature-only model corresponded to the RFC-predicted probability of being physically mature. A gestation adjustment of 1.05 years was applied to the final age estimates (Palmer et al., 2022).

#### iii) Error metric assessment

The accuracy of the resulting epigenetic clocks was tested with Pearson correlation coefficients (*r*) and the calculation of median absolute errors (MAEs) between dental and predicted epigenetic age. We further tested the accuracy of the DNA age predictions by performing a linear regression of DNA predicted age against dental age (Barratclough et al., 2021; Beal et al., 2019; Peters et al., 2023). This analysis allowed us to assess how closely the DNA predicted ages aligned with dental age estimates. A slope close to 1 and an intercept near 0 would indicate a strong agreement between predicted and observed values, while deviations may highlight systematic under- or overestimation. The coefficient of determination (R^2^) was also examined to quantify the proportion of variance in chronological age explained by the epigenetic predictions.

#### iv) Effect of tissue decomposition

To assess whether tissue decomposition influenced the accuracy of epigenetic age predictions, we compared the absolute age prediction errors from the ‘*relaxed*’ subset across decomposition condition categories DCC 1 to 3 (fresh to moderate) assigned post-mortem during sampling (IJsseldijk et al., 2019). Absolute prediction errors were calculated as the absolute difference between predicted and observed ages for each individual. We then used a Kruskal–Wallis rank sum test to test for differences in error distributions among decomposition condition categories, as the error data did not meet the assumptions of normality.

#### v) Effect of storage duration

To evaluate whether long-term storage of samples could bias epigenetic age estimates, storage time was calculated as the difference between the year of DNA methylation analysis (2023) and the year of sample collection, as indicated in the sample identifier. Absolute age prediction errors from across all samples were used as the response variable. Associations between storage time and prediction error were assessed using Spearman rank correlation (robust to deviations from normality) and linear regression, with and without chronological age included as a covariate. For the regression models, residuals were inspected using diagnostic plots and tested for normality using the Shapiro–Wilk test to confirm model assumptions.

#### vi) Age predictions based on other odontocetes clocks

Additionally, our clock’s performance was tested in relation to that of other epigenetic clocks, using the `MammalMethylClock` R package browsing the Clock database with the search criteria odontocetes and skin samples. We applied previously published multi-species and species-specific odontocetes clocks to our samples and compared the respective MAE and *r*-values between the resulting clocks.

### 2.6. Sex prediction

To predict the sex of each sample based on the methylation data, we used the R package `glmnet` following the approach of Peters et al., 2023 using a binomial elastic net regression with α = 0.4 without LOOCV. The sex was encoded as binary outcome variable (0 = female, 1 = male; Peters et al., 2023). With a predicted probably > 0.5, the sample was male (Peters et al., 2023).

## 3. Results

Following Zoller & Horvath (2024), eight outliers collected between 2001 and 2008 were removed as part of the sample quality assessment (e.g., hierarchical clustering, failed probe fractions, and principal component analysis; Supplementary Material, Table S2). In addition to these technical outliers, one further animal (KS19-23Dd) was excluded due to inconsistencies between dental age, pectoral bone age, and number of *corpora albicans*. Accordingly, analyses continued with a total sample size of 75 animals.

### 3.1. Age prediction

#### i) Elastic net regression with LOOCV

The elastic net models with LOOCV showed a strong correlation between dental age and predicted age across the different subsets (Figure 1).

**Figure 1.**
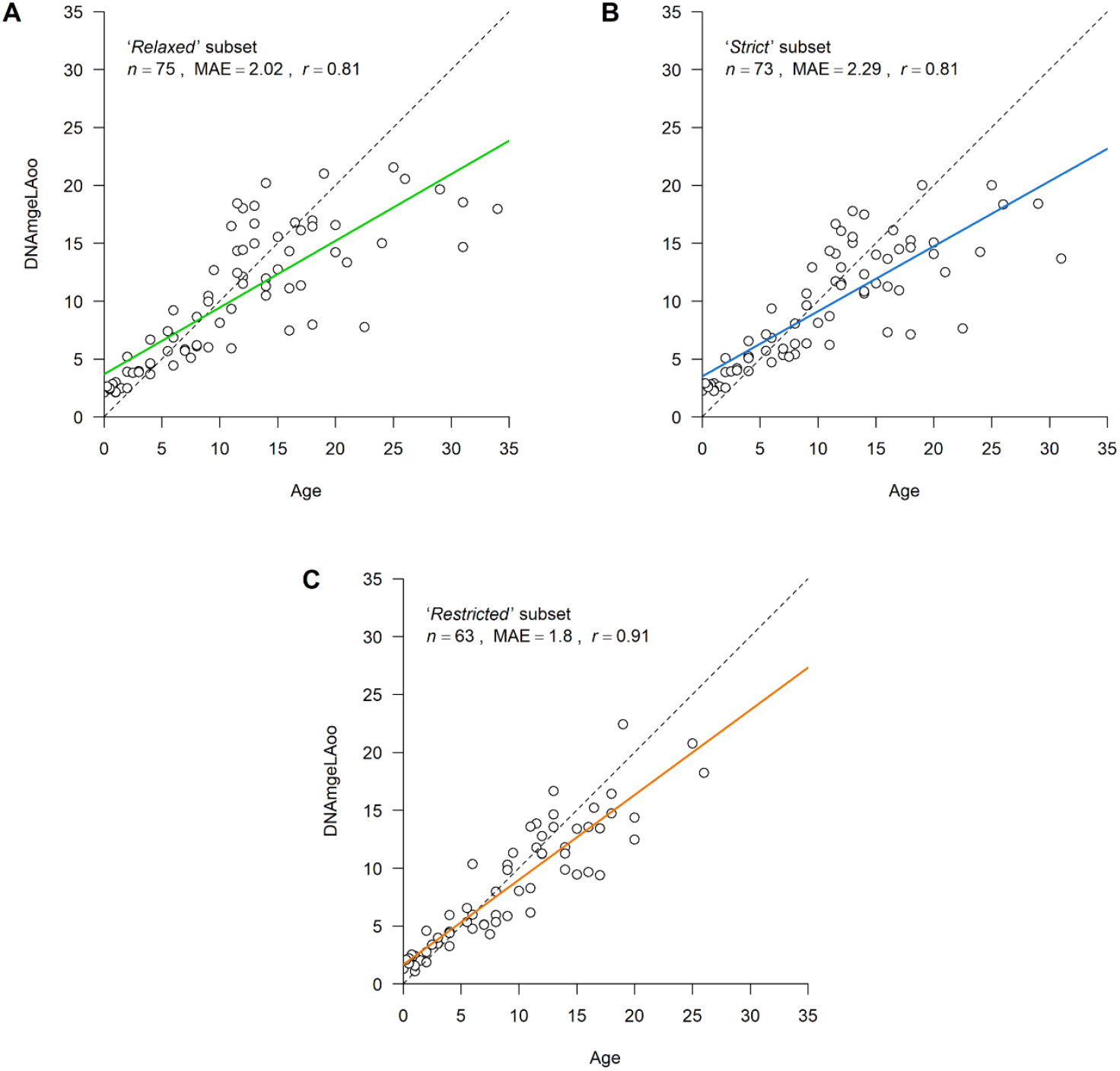
Epigenetic ages (DNAmAgeLOO) estimated using elastic net regression models with Leave-One-Out Cross-Validation (LOOCV) applied to skin samples of common dolphin (*Delphinus delphis*) for the A) ‘*relaxed*’, B) ‘*strict*’, and C) ‘*restricted*’ subsets. The ‘*relaxed*’ subset included all individuals, the ‘*strict*’ subset excluded those with only estimated age ranges or minimum ages, and the ‘*restricted*’ subset further removed outliers with absolute errors > 6 years. Regression lines are shown in green (‘*relaxed*’), blue (‘*strict*’), or orange (‘*restricted*’), with the dotted diagonal indicating perfect correlation (*y* = *x*). Individual animals are shown as dots. Sample size (n), Pearson correlation (*r*), and median absolute error (MAE) are reported for each model.

For the *‘relaxed’* subset, the elastic net regression retained 38 CpG sites (Supplementary Material, Table S3) and produced a MAE of 2.02 years, with a correlation of *r* = 0.81 (*p* = 1.39 × 10^−18^) and an R^2^ of 0.66 using a LOOCV. In 12 individuals, the absolute difference between actual and predicted age exceeded 6 years (Table 1). While predicted age was overestimated in three younger individuals (aged 11.5, 12 and 14 years), most discrepancies (*n* = 9) involved underestimation in older dolphins with a dental age greater than 16 years.

**Table 1:** Error metrics for elastic net regression models predicting epigenetic age in common dolphin (*Delphinus delphis*), evaluated with Leave-One-Out Cross-Validation (LOOCV) for the ‘*relaxed*’, ‘*strict*’, and ‘*restricted*’ subsets. The ‘*relaxed*’ subset included all individuals; the ‘*strict*’ subset excluded those with estimated age ranges or minimum ages; and the ‘*restricted*’ subset further removed outliers with errors > 6 years. Shown are the mean and median absolute errors, the number of samples with errors > 6 and 10 years, and the maximum absolute error.

| Metric | ‘Relaxed’ | ‘Strict’ | ‘Restricted’ |
| --- | --- | --- | --- |
| Mean Absolute Error | 3.42 | 3.22 | 2.21 |
| Median Absolute Error | 2.02 | 2.29 | 1.80 |
| N samples with Absolute Error > 6 years | 12 | 9 | 4 |
| N samples with Absolute Error > 10 years | 5 | 4 | 0 |
| Maximum Absolute Error | 16.31 | 17.33 | 7.77 |

The training set of the *‘strict’* subset retained 24 CpG sites (Supplementary Material, Table S3) and yielded a MAE of 2.29 years, with a correlation of *r* = 0.81 (*p* =2.33 × 10^−18^) and an R^2^ of 0.66 using LOOCV. Nine samples, all from individuals older than 16 years, had an absolute error exceeding 6 years, with the predicted ages consistently underestimated in all cases.

For the *‘restricted’* subset, the elastic net model retained 22 CpG sites (Supplementary Material, Table S3) and demonstrated the best performance, achieving a MAE of 1.80 years, a correlation of *r* = 0.91 (*p* = 1.48 × 10^−24^), and an R^2^ of 0.82 using LOOCV. In this subset, four individuals — each over 16 years of age — had an absolute error exceeding 6 years. Notably, all again were underestimated. A summary of error metrics of the LOOCV are provided in Table 1.

All elastic nets with LOOCV showed the tendency of overestimating age in younger individuals and underestimating age in older animals (‘*relaxed*’: regression slope: 0.58, regression intercept: 3.69; ‘*strict*’: regression slope: 0.56, regression intercept: 3.52; ‘*restricted*’: regression slope: 0.73, regression intercept: 1.69). The *‘restricted’* subset has the slope closest to 1, indicating that it exhibits the least amount of bias among the three, performing best in maintaining proportional predictions across ages.

#### ii) Hybrid epigenetic clock

The hybrid model using the ‘*relaxed*’ subset produced a median absolute error (MAE) of 2.61 years (Supplementary Material, Figure S2) with a corelation of *r* = 0.73 (*p* = 6.16 × 10^−14^). Linear regression of DNA predicted age against dental age yielded a slope of 0.5, an intercept of 4.15, and an R^2^ of 0.54.

#### iii) Effect of tissue decomposition

We found no difference in absolute epigenetic age prediction error across the decomposition condition categories (*p* = 0.746, chi-squared: 1.23, df = 3), indicating fresh to moderate decomposition change had no overall effect on the accuracy of the epigenetic age estimates.

#### iv) Effect storage duration

Storage duration in the ‘relaxed’ subset ranged from 0 to 23 years (median = 9 years). Spearman correlation indicated no association between storage time and prediction error (ρ = 0.10, p = 0.38). Similarly, linear regression showed that storage time did not predict error (β = 0.029 ± 0.064 SE, p = 0.65, R^2^ = 0.003). Residuals from this simple model deviated from normality; however, when chronological age was included as a covariate, residuals conformed to model assumptions (Shapiro– Wilk p = 0.10). In this adjusted model, storage time remained non-significant (β = 0.048 ± 0.045 SE, p = 0.30), while age strongly predicted error (β = 0.316 ± 0.036 SE, p = 7.3 × 10^−13^, R^2^ = 0.51). Together, these results indicate that storage duration did not confound clock performance in our dataset.

#### v) Age predictions based on other odontocetes clocks

Age predictions for our samples, generated using both species-specific and multi-species odontocete clocks from the Clock Database, are shown in Supplementary Material (Figure S3), with corresponding error metrics in Table S4.

The multispecies odontocetes clock (Robeck, Fei, Lu, et al., 2021) predicted the age of our samples with an accuracy of *r* = 0.79 for the ‘*relaxed*’, *r* = 0.8 for the ‘*strict*’ and *r* = 0.85 for the ‘*restricted*’ subset, and a MAE of 11.8, 11.84 and 12.42, respectively. The use of the multispecies odontocete blood and skin clock resulted in comparable MAE values, but the latter revealed lower age correlations (Supplementary Material, Table S4). Comparing our results to that of other species-specific clocks, the Hector’s and Māui dolphin (*Cephalorhynchus hectori hectori* and *C. h. maui*; Hernandez et al., 2023) clock performed best for our samples regarding the MAE between actual and predicted age, with a MAE ranging from 2.71 to 3.01 depending on the subset of samples used (Supplementary Material, Table S4). While both multispecies clocks showed the tendency of overestimating the age of our samples, the *Cephalorhynchus* clock consistently underestimated samples above the age of 6 years (see Supplementary Material; Text S1 and S2).

### 3.2. Sex prediction

The elastic net model retained 197 CpG sites and achieved 100% accuracy in predicting sex. The probability of males being correctly classified as male ranged from 0.9892 to 0.9953, while the probability of females being misclassified as male was very low, ranging from 0.0025 to 0.0078.

## 4. Discussion

Here we present the first species-specific epigenetic clock for common dolphins, developed using DNA methylation profiles from skin samples of individuals with estimated dental ages. The elastic net regressions with LOOCV exhibited robust performance across all three models, with the ‘*restricted*’ subset (MAE = 1.80, *r* = 0.91, R^2^ = 0.82) yielding the highest accuracy with a MAE equivalent to approximately ± 5.1% of maximum life expectancy (MLE: 35 years, Murphy et al., 2014). The ‘*relaxed*’ and ‘*strict*’ models also performed strongly, with MAEs (2.02 and 2.29 years) equivalent to approx. ± 5.8% and ± 6.5% of maximum life expectancy, and correlation coefficients of 0.81.

Epigenetic clocks have been shown to estimate chronological age with high accuracy (Jylhävä et al., 2017). However, even when trained on extensive datasets encompassing all age groups, these clocks still exhibit variability among individuals, which could reflect differences in their biological aging rates (Jylhävä et al., 2017). Variation in aging rates might be influenced by numerous factors, including reproductive history (Shirazi et al., 2020), early-life adversities (Colich et al., 2020), nutrition (Fitzgerald et al., 2021; Weindruch et al., 2001), and exposure to environmental contaminants (Liu et al., 2021). In cetaceans, eleven odontocete epigenetic clocks have been developed to date, using a range of sample types including skin, blood, multi-tissue, and faeces (Supplementary Material, Table S5). Notably, only three prior studies calibrated their epigenetic clocks using growth layer groups from stranded and bycaught animals, with concerns raised that reliance on tooth age calibration may compromise the robustness of epigenetic clocks (Bors et al., 2021; Hernandez et al., 2023; Mori et al., 2024; Zoller et al., 2025).

While dental aging provides robust age estimates for animals younger than 15 years, the reliability of this method decreases in older individuals (Barratclough et al., 2023; Betty et al., 2023; Murphy et al., 2014). Specifically, factors such as tooth wear, the compression of growth layer groups, and the presence of accessory lines, can compromise the accuracy of age estimates (Barratclough et al., 2023; Betty et al., 2023; Murphy et al., 2014). Such factors generally result in dental aging underestimation of older individuals (Barratclough et al., 2023), especially when aging is restricted to a single tooth, as is typical in studies of animals under human care (Barratclough et al., 2023). To mitigate this, we processed up to 3 teeth per animal, further selected the least worn and straightest teeth, and assigned minimum age estimates when uncertainty remained due to accessory lines or compressed growth layer groups. Despite these precautions, our epigenetic clock repeatedly underestimated the ages of older individuals (i.e., >16 years) across all subsets.

The consistent underestimation of older dolphins suggests that discrepancies between dental and epigenetic age assessments unlikely stem from inaccuracies in the dental aging method itself. Specifically, if dental aging inaccuracies were the primary source of error, we would expect epigenetic age estimate to exceed dental ages in older individuals — which was not observed. Additionally, contrary to our first hypothesis, overall performance was similar across both the ‘*relaxed*’ and ‘*strict*’ subsets. Notably, the ‘*relaxed*’ model yielded a lower MAE and a regression slope closer marginally closer to 1, indicating more balanced age predictions across the lifespan. This result suggests that excluding samples with uncertain dental ages did not substantially improve model performance. However, the performance of the ‘*relaxed*’ model was comparable to most previously published skin-based odontocete clocks calibrated using growth layer groups, though fell marginally short of the accuracy reported for clocks developed from known-age individuals in long-term observational studies or human care (Supplementary Material, Text S2). We considered whether this performance gap could be explained by factors relating to tissue origin. Our samples originated from a variety of contexts, including mass and single strandings, bycatch events, and one individual under human care. While mass strandings are typically unrelated to individual health problems (Cordes, 1982), single strandings may involve animals affected by illness, compromised health, or poor nutritional status (Arbelo et al., 2013; Boys, 2022). Although poor health has been linked to accelerated epigenetic aging in other species (Newediuk et al., 2025), this contrasts with the consistent underestimation observed here in older individuals.

We also considered whether post-mortem decomposition or storage duration could influence methylation patterns. While post-mortem changes may potentially affect methylation at specific loci, a comparison of live and deceased human samples showed comparable methylation-based age predictions (Dias et al., 2020). Consistent with this, we found no significant effect of decomposition condition on epigenetic age prediction error, suggesting that our clock is likely applicable to both live and deceased animals. Similarly, the duration of storage showed no significant association with prediction error i.e., long-term storage did not affect clock performance.

Beyond biological and technical factors, the composition of the training dataset itself plays a critical role in shaping clock performance. While our study encompassed the full estimated lifespan of common dolphins, most samples were from individuals aged 0–20 years. Skin samples from older common dolphins remain rare and the limited and fragmentary representation of older animals may have reduced the accuracy of age estimations in the upper age ranges. When designing epigenetic clocks for wildlife, it is crucial to minimise where possible, age-related sampling biases (Newediuk et al., 2025). Specifically, to ensure accurate age predictions, samples should be evenly distributed across age classes, particularly from both young and aged individuals. This prevents poor performance of clocks due to narrow or skewed age distributions (Newediuk et al., 2025). However, heterogenous age distribution of stranded and bycaught individuals relies upon extensive tissue archives which capture the full age spectrum of the species.

In our study, regression analyses revealed slopes smaller than 1 for all three models, indicating a systematic overestimation of age in younger individuals and underestimation in older individuals. This pattern is consistent across multiple species (Beal et al., 2019; Bors et al., 2021; Peters et al., 2023; Polanowski et al., 2014). Among our models, the ‘*restricted*’ model showed the slope closest to 1, suggesting the most balanced age predictions across the lifespan. The phenomenon of ‘regression to the mean’ poses a well-recognized challenge for epigenetic clock models, particularly when predicting ages beyond the range of the training data. This statistical effect causes extreme values to shift toward the population mean, leading to underestimation of older individuals and overestimation of younger ones when these age groups are underrepresented in the training set. In human studies, this has prompted efforts to develop specialised clocks for centenarians (Dec et al., 2023). Indeed, this concept and its implications for research design, have been widely discussed (Campbell & Kenny, 1999). Accordingly, when interpreting epigenetic age estimates for individuals at the extremes of the age spectrum, it is important to account for the influence of regression to the mean and its bias toward the average age of the training cohort.

Regression to the mean, due to sparse representation of older individuals, is one explanation for the underestimation of age at higher values. Another plausible factor is the biological deceleration of DNA methylation change in later life. For example, human epigenetic clocks have been shown to progress rapidly during development and then decelerate, leading to systematic underestimation of age in older individuals (Horvath, 2013b; Kuzub et al., 2022; Marioni et al., 2019). If odontocetes exhibit a comparable biological deceleration, it would systematically bias clock estimates downward in older dolphins, regardless of sample distribution. This has also been hypothesized in bottlenose dolphins, where the use of hybrid models has been suggested as a strategy to address both uneven age distributions and life stage–dependent variation in methylation change (Barratclough et al., 2024).

Hybrid models aim to improve predictions across the lifespan by adjusting for shifts in methylation patterns after physical maturity. Barratclough et al., (2024) applied this strategy in bottlenose dolphins (*Tursiops truncatus*), combining a random forest classifier with two elastic net regression models to improve age prediction for older individuals. However, applying a similar approach to common dolphins yielded a poorer fit than elastic net regression alone, likely due to the small number of physically mature individuals in our dataset (*n* = 14), limiting the classifier’s and mature-age model’s effectiveness. Future approaches such as learning curves (Perlich et al., 2003) or bias–variance decomposition (Domingos, 2000), could be used to disentangle whether underperformance reflects model bias or limited sample size when sufficient sample sizes permit.

Despite limitations identified, our results highlight the promise of epigenetic clocks based on dental aging in addressing long-standing challenges in age estimation, particularly for species where invasive sampling or long-term observational data are not feasible. Beyond age estimation, our clock also accurately predicts sex based on methylation patterns at CpG sites, achieving 100% accuracy, a finding consistent with previous studies (Bors et al., 2021; Peters et al., 2023; Robeck, Fei, Haghani, et al., 2021). Tools that enable accurate age and sex estimation are becoming increasingly critical given the significant anthropogenic threats facing common dolphins globally (Abraham et al., 2021; Bilgmann et al., 2014; Fernández-Contreras et al., 2010; Hamer et al., 2007; Mannocci et al., 2012; Murphy et al., 2013; Mussi et al., 2021; Peltier et al., 2016, Peltier et al., 2021; Piroddi et al., 2011; Tulloch et al., 2020; Vella et al., 2021), where several populations are in decline (Bearzi et al., 2008; Murphy et al., 2013; Piroddi et al., 2011). Understanding life-history parameters is critical for conservation management (Betty et al., 2019; Heydenrych et al., 2021; Manlik et al., 2022). For example, accurate age estimates underpin population viability analyses and are essential to quantify population-level impacts such as fisheries bycatch (Manlik et al., 2016; Palmer et al., 2022; Verborgh et al., 2021). Since post-mortem age assessment may bias survivorship estimates (Betty et al., 2023) used to evaluate fisheries impacts, our epigenetic clock provides a critical advance by enabling age estimation from living animals.

The ability to age live individuals strengthens conservation efforts globally. Life-history traits are comparable among common dolphin populations worldwide (Danil & Chivers, 2007; Murphy & Rogan, 2006; Palmer et al., 2022, 2023; Westgate & Read, 2007), and species-specific epigenetic clocks have been shown to accurately predict age in subspecies and closely related sister taxa (Barratclough et al., 2024, Barratclough et al., 2025; Peters et al., 2023; Robeck, Fei, Lu, et al., 2021), our epigenetic clock has broad applicability for this cosmopolitan oceanic species. Our clock further may facilitate comparisons of epigenetic age acceleration across populations experiencing varying environmental pressures (Newediuk et al., 2025). By enabling reliable age and sex estimation, this tool can enhance conservation strategies and improve our understanding of population dynamics, both in Aotearoa New Zealand and throughout the species’ range. Finally, with concerns of decomposition and tooth aging effects on clock calibration addressed here, future methylated clocks should be considered for stranded and bycaught dolphin populations, especially those underpinned by longitudinal tissue archives that allow teeth estimates to be further cross validated with wider life history data.

## 7. Data availability

All data and R code supporting this study are accessible at https://github.com/Ehanninger/Epigenetic-clock-Delphinus-delphis. The dataset will further be accessible through the Mammalian Methylation Consortium’s data release. The mammalian methylation array used in this research is available from the non-profit Epigenetic Clock Development Foundation.

## 8. Author contributions

EMFH performed conceptualization, data curation, formal analysis, funding acquisition, investigation, methodology and draft writing. KJP supported with conceptualization, formal analysis, investigation and methodology, draft writing and supervision. LG supported with conceptualization, methodology, formal analysis, and manuscript drafting; AB assisted in methodology, formal analysis and draft writing; ELB conducted methodology and supported in manuscript drafting; EIP conducted methodology and supported in manuscript drafting. SH supported in manuscript drafting. KAS performed conceptualization, data curation, funding acquisition, methodology, draft writing and supervision.

## 9. Acknowledgments

This work was funded by Massey University Research Fund (RM26712 awarded to KAS) and Massey University Wildbase Research Trust Fund (RM25735 awarded to EH). Permits for sample collection (39239-MAR & 111522-MAR) and sample export (111445-MAR) were obtained from the New Zealand Department of Conservation Te Papa Atawhai. We are particular grateful to local iwi and hapū (Indigenous New Zealanders) for supporting scientific data and sample collection and their approval to access their taonga for this study. We also acknowledge Max Clark, Hannah Hendriks and Marlane Felsing (Department of Conservation Te Papa Atawhai) for their assistance with iwi consultation, permitting, Robert Brooke (Epigenetic Clock Development Foundation) and Razvan Hancu (New Zealand Ministry for Primary Industries) for their logistical support. Access to samples was possible thanks to members of the Cetacean Ecology Research Group, with histological support from Matthew Perrott, Evelyn Lupton and Petru Daniels (Massey University).

## Footnotes

1 https://genomics.senescence.info/species/index.html

