## Supplementary Materials for "Dental aging offers new insights to the first epigenetic clock for common dolphins (*Delphinus delphis*)"

### Supplementary text

#### Text S1 Extended results

##### Age predictions based on other odontocetes clocks

Age predictions for our samples, generated using both multi-species and species-specific odontocetes clocks from the Clock database, are presented in Figure S3 and Table S4.

The multispecies odontocetes clock (Robeck, Fei, Lu, et al., 2021) predicted the age of our samples with an accuracy of $r$ = 0.79 for the '*relaxed*', $r$ = 0.8 for the '*strict*' and $r$ = 0.85 for the '*restricted*' subset, and a MAE of 11.8, 11.84 and 12.53, respectively. The use of the multispecies odontocete blood and skin clock resulted in comparable MAE values, but the latter revealed lower age correlations (Supplementary Material, Table S4). Comparing our results to that of other species-specific clocks, the *Cephalorhynchus* clock (*C. hectori hectori* and *C. h. maui*; Hernandez et al., 2023) clock performed best for our samples regarding the MAE between actual and predicted age, with a MAE ranging from 2.71 to 3.01 depending on the subset of samples used (Supplementary Material, Table S4). While both multispecies clocks showed the tendency of overestimating the age of our samples, the *Cephalorhynchus* clock consistently underestimated samples above the age of 6 years (see Supplementary Material; Text S1 and S2).

### Text S2 Extended discussion

#### Age predictions based on other odontocetes clocks

For the '*relaxed*' subset, the skin and the skin/ blood multi-species odontocetes clock resulted in a similar MAE of 11.8 and 11.54 years, respectively (Figure S3 & Table S4). However, the skin clock ($r$ = 0.79) demonstrated a better correlation between dental and predicted age, compared to the skin and blood clock ($r$ = 0.68). This is surprising since methylation rates are highly conserved in blood and blood-based clocks are considered more precise (Robeck, Fei, Haghani, et al., 2021; Robeck, Fei, Lu, et al., 2021). Age prediction would have been expected to improve when applying the combined skin and blood clock to our samples. However, similar observations have been previously reported, and it was assumed that the differences between the skin epitome of different species might be larger than differences in the blood epitome (Peters et al., 2023). Both, the skin and the skin and blood multi-species odontocetes clocks showed the tendency of overestimating the age of our samples. While samples of common dolphins were used in the construction of the multispecies odontocete clock, the sample size was limited (skin samples: $n$ = 3, blood samples: $n$ = 3) and samples of longer-lived odontocetes, such as bottlenose dolphins (*Tursiops truncatus*) were more strongly represented (skin samples: $n$ = 41, blood samples: $n$ = 140; Robeck, Fei, Lu, et al., 2021). This might explain the overestimation of age in our samples due to differences in life-history features and maximum lifespan. Our findings align with previous work (Barratclough et al., 2021; Peters et al., 2023) suggesting that the accuracy of epigenetic age estimation can be enhanced through the development of species-specific clocks.

When testing other available odontocetes clocks, the epigenetic clock for *Cephalorhynchus* (*C. hectori hectori* and *C. h. maui*, Hernandez et al., 2023) performed best in estimating the age of our samples with a MAE of 3.01 for the '*relaxed*' subset and a correlation of $r$ = 0.79. However, the clock consistently underestimated the age of our samples for individuals with an age > 6 years. For Hector's dolphins, there is a lack of knowledge on life-history features, such as age at sexual maturity and reproductive rates (Hernandez et al., 2023). While currently unknown, lifespan has been estimated to be around 20 years (Hernandez et al., 2023; Slooten & Lad, 1991). Given that the *Cephalorhynchus* clock is calibrated for a faster-aging species, this may account for the underestimation observed in our common dolphin samples, which are believed to have a maximum lifespan of around 30 years (Perrin, 2009). While age at attainment of sexual maturity of Hector's dolphins is currently unknown (Hernandez et al., 2023), our results suggest a similar aging trajectory of Hector's and common dolphins up until the age of 6. The similarity in epigenetic age estimates between Hector's and common dolphin up to age 6 may reflect conserved early-life developmental processes, with divergence emerging later due to differences in timing of maturity and species-specific aging trajectories.

#### Error metrics reported by previous literature

Error metrics reported by previously published odontocetes clocks are shown in Table S5. These clocks were developed using a range of tissue types, including skin, blood, multi-tissue, and faeces; however, our comparison focuses specifically on skin-based clocks. Clocks calibrated using growth layer groups (Bors et al., 2021; Hernandez et al., 2023; Mori et al., 2024) report MAEs equivalent to approximately ± 10.45% of maximum life expectancy (MLE) for Hector's and Māui dolphins (MLE: 20 years; Slooten, 1991), ±5% and for beluga whales (MLE: 60–70 years; Suydam, 2009). The predictive performance of these clocks is reflected in a Pearson correlation coefficient of $r$ = 0.87 for Hector's and Māui dolphins (Hernandez et al., 2023) and a coefficient of determination of $R^2$ = 0.74 for beluga whales (Bors et al., 2021). For Risso's dolphins no median absolute error was reported, but a $R^2$ of 0.71 (Mori et al., 2024). Clocks developed from long-term observational data show lower relative errors: Peters et al., (2023) reported an MAE of ± 3.13–4.2% ($r$ = 0.86) for Indo-Pacific bottlenose dolphin (MLE: 40–50 years; Wang & Yang, 2009), while Parsons et al., (2023) reported ± 2.25% ($r$ = 0.96) for killer whales (MLE: up to 90 years; Foote, 2008). In another observational study of bottlenose dolphins, Beal et al., (2019) reported an $R^2$ of 0.78, though no MAE was provided. A clock developed using samples from animals in human care achieved an MAE of ± 4.9% of maximum life expectancy for bottlenose dolphins (MLE: 51.6 years; Weigl, 2005) and a correlation of $r$ = 0.95 (Robeck, Fei, Haghani, et al., 2021).

#### User guide and limitations

This epigenetic clock provides a robust framework for age estimation in common dolphins (*Delphinus delphis*). It shows highest accuracy within the age range represented by the majority of our training data (up to ~16 years), while estimates for older individuals remain informative but increasingly underestimate chronological age. This underestimation likely reflects both sparse representation of older dolphins in the calibration dataset and a biological slowing of methylation changes with age. Neither decomposition state nor storage duration influenced prediction accuracy, indicating that these factors did not confound our dataset and that the clock is likely applicable to live animals. Although the calibration dataset was derived from New Zealand dolphins, the broader utility of this clock is supported by the demonstrated reliability of epigenetic clocks in other species, which have proven consistent across populations and even transferable among closely related taxa.

#### Supplementary tables

**Table S1:** Common dolphin (*Delphinus delphis*) sample IDs included in the '*relaxed*', '*strict*', and '*restricted*' subsets used for epigenetic clock model development. For each sample, the dental age and sex are provided, as well as the decomposition condition category (DCC), indicating carcass preservation at the time of sampling (IJsseldijk et al., 2019).

| ID | DCC | Dental Age | Sex | Subset<br>'Relaxed' | <br>'Strict' | <br>'Restricted' |
|---|---|---|---|---|---|---|
| WB00-06Dd | 2 | 6 | F | ✓ | ✓ | ✓ |
| WB02-01Dd | 2 | 19 | M | ✓ | ✓ | ✓ |
| WS02-38Dd | 2 | 20 | F | ✓ | ✓ | ✓ |
| WS02-40Dd | N/A | 13 | M | ✓ | ✓ | ✓ |
| WB03-18Dd | 2 | 8 | M | ✓ | ✓ | ✓ |
| WS03-42Dd | 2-3 | 25 | F | ✓ | ✓ | ✓ |
| WS03-43Dd | 2 | 13 | F | ✓ | ✓ | ✓ |
| WB04-25Dd | 3 | 9 | F | ✓ | ✓ | ✓ |
| WS04-35Dd | 2 | 15 | F | ✓ | ✓ | ✓ |
| WS05-25Dd | 2 | 7 | F | ✓ | ✓ | ✓ |
| WS06-09Dd | 2 | 14 | F | ✓ | ✓ | ✓ |
| KS07-12Dd | 1 | 16 | F | ✓ | ✓ | ✓ |
| KS09-08Dd | 2 | 1 | F | ✓ | ✓ | ✓ |
| KS09-11Dd | 2 | 3 | F | ✓ | ✓ | ✓ |
| KS09-13Dd | 1 | 8 | M | ✓ | ✓ | ✓ |
| KS09-18Dd | 2 | 11 | F | ✓ | ✓ | ✓ |
| KS09-29Dd | 2 | 0 | M | ✓ | ✓ | ✓ |
| KS10-01Dd | 2 | 2 | F | ✓ | ✓ | ✓ |
| KS10-09Dd | 2-3 | 4 | F | ✓ | ✓ | ✓ |
| KS10-15Dd | 2 | 0.75 | M | ✓ | ✓ | ✓ |
| KS10-27Dd | 3 | 4 | M | ✓ | ✓ | ✓ |
| KS10-29Dd | 2 | 1 | M | ✓ | ✓ | ✓ |
| KS11-08Dd | 1 | 6 | M | ✓ | ✓ | ✓ |
| KS11-12Dd | 3 | 6 | M | ✓ | ✓ | ✓ |
| KS11-13Dd | 2 | 17 | M | ✓ | ✓ | ✓ |
| KS11-27Dd | 1 | 2 | M | ✓ | ✓ | ✓ |
| KS11-39Dd | 2 | 2 | F | ✓ | ✓ | ✓ |
| KS12-14Dd | 2 | 11 | M | ✓ | ✓ | ✓ |
| KS12-17Dd | 2 | 1.5 | M | ✓ | ✓ | ✓ |
| KS12-23Dd | 2 | 2.5 | M | ✓ | ✓ | ✓ |
| KS13-09Dd | 2 | 5.5 | M | ✓ | ✓ | ✓ |
| KS14-38Dd | 2 | 16 | M | ✓ | ✓ | ✓ |
| KS14-42Dd | 2 | 1 | F | ✓ | ✓ | ✓ |
| KS14-55Dd | 3 | 3 | F | ✓ | ✓ | ✓ |
| KS14-56Dd | 3 | 18 | F | ✓ | ✓ | ✓ |
| KS14-57Dd | 3 | 14 | F | ✓ | ✓ | ✓ |
| KS15-16Dd | 3 | 26 | M | ✓ | ✓ | ✓ |
| KS15-20Dd | 2 | 0.5 | F | ✓ | ✓ | ✓ |
| KS15-21Dd | 2 | 4 | F | ✓ | ✓ | ✓ |
| KS16-06Dd | 2-3 | 0.5 | F | ✓ | ✓ | ✓ |

**Table S1:** Continued.

| ID | DCC | Dental Age | Sex | Subset 'Relaxed' | Subset 'Strict' | Subset 'Restricted' |
|---|---|---|---|---|---|---|
| KS16-27Dd | 3 | 7.5 | M | ✓ | ✓ | ✓ |
| KS17-02Dd | 2 | 11.5 | M | ✓ | ✓ | ✓ |
| KS17-08Dd | 2 | 4 | M | ✓ | ✓ | ✓ |
| KS19-10Dd | 2-3 | 7 | F | ✓ | ✓ | ✓ |
| KS19-12Dd | 2-3 | 9 | M | ✓ | ✓ | ✓ |
| KS19-13Dd | 2-3 | 5.5 | F | ✓ | ✓ | ✓ |
| KS19-14Dd | 3 | 10 | F | ✓ | ✓ | ✓ |
| KS19-17Dd | 2-3 | 12 | F | ✓ | ✓ | ✓ |
| KS19-18Dd | 2 | 0.25 | M | ✓ | ✓ | ✓ |
| KS19-19Dd | 2 | 11 | F | ✓ | ✓ | ✓ |
| KS19-20Dd | 2 | 12 | F | ✓ | ✓ | ✓ |
| KS19-21Dd | 2-3 | 15 | F | ✓ | ✓ | ✓ |
| KS19-22Dd | 2 | 18 | F | ✓ | ✓ | ✓ |
| KS19-39Dd | 2-3 | 8 | F | ✓ | ✓ | ✓ |
| KS22-03Dd | 2 | 9 | F | ✓ | ✓ | ✓ |
| KS23-05Dd | 1 | 20 | F | ✓ | ✓ | ✓ |
| KS23-07Dd | 1 | 13 | F | ✓ | ✓ | ✓ |
| KS23-10Dd | 2 | 14 | F | ✓ | ✓ | ✓ |
| KS23-11Dd | 2 | 17 | F | ✓ | ✓ | ✓ |
| KS23-43Dd | 1 | 16.5 | M | ✓ | ✓ | ✓ |
| KS23-44Dd | 2 | 9.5 | M | ✓ | ✓ | ✓ |
| KS23-47Dd | 2 | 11.5 | M | ✓ | ✓ | ✓ |
| KS23-50Dd | 2 | 12 | M | ✓ | ✓ | ✓ |
| WS04-36Dd | 2 | 22.5 | F | ✓ | ✓ | X |
| W08-17Dd | 1 | 29 | F | ✓ | ✓ | X |
| KS09-14Dd | 3 | 18 | F | ✓ | ✓ | X |
| KS10-06Dd | 2-3 | 16 | F | ✓ | ✓ | X |
| KS14-51Dd | 3 | 14 | M | ✓ | ✓ | X |
| KS17-01Dd | 2 | 12 | M | ✓ | ✓ | X |
| KS22-52Dd | 3 | 11.5 | M | ✓ | ✓ | X |
| KS23-42Dd | 1 | 21 | M | ✓ | ✓ | X |
| KS23-45Dd | 2 | 31 | M | ✓ | ✓ | X |
| KS23-46Dd | 1 | 24 | M | ✓ | ✓ | X |
| W08-16Dd | 2 | ≥ 34 | F | ✓ | X | X |
| KS14-54Dd | 3 | ≥ 31 | F | ✓ | X | X |

**Table S2:** Sample quality control procedures applied to ensure data integrity for downstream DNA methylation analysis in common dolphins (*Delphinus delphis*). Samples were excluded based on probe detection p-values (>1% failed probes; fraction outliers), principal component analysis (PCA), and hierarchical clustering (HC), which were performed to identify technical outliers and samples with aberrant DNA methylation profiles. Additionally, sample KS19-23Dd was excluded as a life-history anomaly based on pectoral fin radiography and ovarian corpora albicans counts. DCC refers to the tissue decomposition condition category indicating carcass preservation at the time of sampling (IJsseldijk et al., 2019).

| ID | DCC | PCA Outlier | Fraction Outlier | HC Outlier | Life-history outlier |
|---|---|---|---|---|---|
| WS06-05Dd | N/A | Yes | - | - | - |
| WS02-03Dd | 2 | Yes | - | - | - |
| WB01-43Dd | 3 | Yes | - | - | - |
| WB04-04Dd | N/A | Yes | - | - | - |
| WS05-37Dd | 2 | Yes | Yes | Yes | - |
| WS06-15Dd | 2 | - | Yes | - | - |
| KS08-02Dd | 3 | Yes | Yes | Yes | - |
| KS18-07Dd | 2 | Yes | - | - | - |
| KS19-23Dd | 2-3 | - | - | - | Yes |

**Table S3:** Cytosine-phosphate-guanine (CpG) sites retained by the elastic net regression models for estimating epigenetic age in common dolphins (*Delphinus delphis*) for the '*relaxed*', '*strict*', and '*restricted*' subsets. The subsets included different samples, as detailed in Table S1.

| CpG site | Subset | | |
|---|---|---|---|
| | 'Relaxed' | 'Strict' | 'Restricted' |
| Intercept | 2.707102179 | 1.776196521 | 2.187039858 |
| Probe-ID | Model coefficient | Model coefficient | Model coefficient |
| cg00008998 | -0.174793681 | -0.16416 | -0.30144 |
| cg01167424 | -0.039862449 | - | -0.01709 |
| cg01814115 | -0.394351964 | -0.60808 | -0.7008 |
| cg02909927 | -0.015346173 | | - |
| cg03738429 | -0.391731339 | -0.2613 | - |
| cg04010581 | -0.035275789 | -0.10929 | -0.10229 |
| cg04051518 | -0.130802155 | - | -0.21781 |
| cg04905851 | -0.06589915 | - | - |
| cg06796713 | - | - | -0.34456 |
| cg07804445 | -0.592141011 | -0.55117 | -0.46492 |
| cg08377921 | - | - | - |
| cg08550913 | - | - | -1.97353 |
| cg08611749 | -0.158115072 | -0.00516 | - |
| cg09058643 | -0.167154015 | - | - |
| cg09363187 | - | 0.176106 | - |
| cg09461098 | -0.680099807 | -0.81789 | -1.52703 |
| cg09916017 | 0.235893107 | - | - |
| cg10277282 | 0.040840017 | - | - |
| cg11116727 | -0.333141023 | -0.38212 | -0.11971 |
| cg11260459 | - | - | -0.36367 |
| cg12053353 | -0.229100545 | -0.26633 | - |
| cg12373771 | - | - | 0.021864 |
| cg12584622 | -0.026700007 | -0.01643 | - |
| cg13060540 | -0.170448176 | -0.13771 | - |
| cg13260846 | 0.036436683 | - | 0.069161 |
| cg13325480 | - | - | -0.21954 |
| cg14142471 | - | - | -0.20964 |
| cg15303324 | -0.119960222 | -0.19948 | - |
| cg17037716 | -0.011535008 | - | - |
| cg17696987 | -0.014918802 | - | - |
| cg18271715 | - | - | -0.26503 |
| cg18583520 | -1.1124165 | -0.64806 | - |
| cg18734357 | - | - | -0.01144 |
| cg19355776 | - | - | -0.0793 |
| cg19591642 | -0.080327164 | - | - |
| cg20490705 | - | - | -0.01144 |
| cg20582188 | -0.534397174 | -0.56815 | -0.70342 |
| cg21454760 | -0.125015485 | -0.21864 | - |
| cg21898068 | -0.072178198 | -0.08277 | - |
| cg22189866 | -0.344160257 | - | - |
| cg22423049 | -0.052793821 | - | - |
| cg22572209 | -0.462827113 | -0.22744 | - |
| cg23170538 | -0.274028766 | -0.01244 | - |
| cg24888049 | -0.168501987 | - | - |
| cg25184118 | -0.090402011 | - | - |

Table S3: Continued

| | Subset | | |
|---|---|---|---|
| CpG site | 'Relaxed' | 'Strict' | 'Restricted' |
| Probe-ID | Model coefficient | Model coefficient | Model coefficient |
| cg25254739 | -0.399436757 | -0.48822 | -0.33018 |
| cg25402449 | - | -0.05742 | - |
| cg26186239 | 0.002794655 | - | - |
| cg26631126 | -0.552342767 | -0.76605 | -0.78668 |
| cg27038395 | 0.080454226 | 0.276721 | - |
| cg27395516 | -0.548172671 | -0.27504 | - |

**Table S4:** Error metrics for age predictions of common dolphin (*Delphinus delphis*) samples using previously published odontocete epigenetic clocks. Metrics are reported separately for models trained on skin (S) and blood (B) tissue where applicable. Performance is summarised as median absolute error (MAE) and correlation coefficient (*r*) providing an assessment of how accurately these external models predict age in our dataset. The subsets included different samples, as detailed in Table S1.

| | | | 'Relaxed' | | 'Strict' | | 'Restricted' | |
|---|---|---|---|---|---|---|---|---|
| Epigenetic Clock Name | Reference | Tissue | MAE | r | MAE | r | MAE | r |
| Coef.OdontoceteSkinAge.LogLinear2 | Robeck, Fei, Lu, et al., 2021 | S | 11.8 | 0.79 | 11.84 | 0.8 | 12.53 | 0.85 |
| Coef.OdontoceteBloodSkinAge.LogLinear2 | Robeck, Fei, Lu, et al., 2021 | B/S | 11.54 | 0.68 | 11.67 | 0.69 | 12.46 | 0.73 |
| Coef.CetaceanCephalorhynchusHectoriHectoriSkinAge.LogLinear2 | Hernandez et al., 2023 | S | 3.01 | 0.79 | 2.91 | 0.79 | 2.71 | 0.87 |
| Coef.CetaceanDelphinapterusLeucasSkinAge.LogLinear2 | Bors et al., 2021 | S | 7.09 | 0.77 | 7.05 | 0.79 | 6.62 | 0.86 |
| Coef.Killerwhale.Skin | Parsons et al., 2023 | S | 25.09 | 0.77 | 25.41 | 0.78 | 25.41 | 0.84 |
| Coef.IndoPacificBottlenoseSkinClock | Peters et al., 2023 | S | 4.09 | 0.67 | 4.03 | 0.7 | 3.71 | 0.74 |
| Coef.BottlenoseSkin | Barratclough et al., 2021 | S | 48.39 | 0.21 | 48.43 | 0.18 | 48.68 | 0.18 |
| Coef.Bottlenose | Barratclough et al., 2021 | B/S | 27.12 | 0.6 | 27.19 | 0.62 | 27.45 | 0.67 |
| Coef.BottlenoseSkinAge.LogLinear2 | Robeck, Fei, Haghani, et al., 2021 | S | 5.32 | 0.68 | 5.23 | 0.69 | 4.75 | 0.77 |
| Coef.BottlenoseBloodSkinAge.LogLinear2 | Robeck, Fei, Haghani, et al., 2021 | B/S | 31.03 | 0.64 | 31.25 | 0.66 | 31.32 | 0.71 |

**Table S5:** Error metrics reported in published studies for species-specific odontocete epigenetic clocks. Reported metrics include median absolute error (MAE, in years), Pearson's correlation coefficient (*r*), residual error, and coefficient of determination ($R^2$), reflecting the accuracy of chronological age predictions in the respective datasets. Sample size (n) and the tissue type used for model calibration (skin or blood) are also listed for context. These values provide a comparative basis for evaluating the performance of the common dolphin (*Delphinus delphis*) clock developed in this study.

| Species | Tissue | Method of calibration | n | MAE | Residual error | *r* | $R^2$ | Reference |
|---|---|---|---|---|---|---|---|---|
| *C. h. hectori* | Skin | GLGs | 48 | 2.09 | - | 0.87 | - | Hernandez et al., 2023 |
| *D. leucas* | Skin | GLGs | 67 | 2.9 | - | - | 0.74 | Bors et al., 2021 |
| *G. griseus* | Skin | GLGs | 30 | - | - | - | 0.71 | Mori et al., 2024 |
| *T. aduncus* | Skin | Observational | 65 | 2.1 | - | 0.86 | - | Peters et al., 2023 |
| *T. truncatus* | Skin | Observational | 39 | - | 4.83 | - | 0.78 | Beal et al., 2019 |
| *T. truncatus* | Skin | In human care | 87 | 2.53 | - | 0.95 | - | Robeck, Fei, Haghani, et al., 2021 |
| *O. orca* | Skin | Observational | 131 | 2.26 | - | 0.96 | - | Parsons et al., 2023 |
| *T. truncatus* | Blood | In human care | 140 | 1.46 | - | 0.97 | - | Robeck, Fei, Haghani, et al., 2021 |
| *T. truncatus* | Blood/Multi-tissue | In human care | 110 | 2.5 | - | - | 0.95 | Barratclough et al., 2021 |
| *T. truncatus* | Multi-tissue | In human care | 227 | 2.13 | - | 0.93 | - | Robeck, Fei, Haghani, et al., 2021 |
| *T. aduncus* | Faeces | Observational | 36 | 5.08 | - | - | 0.33 | Yagi et al., 2023 |

### Supplementary figures

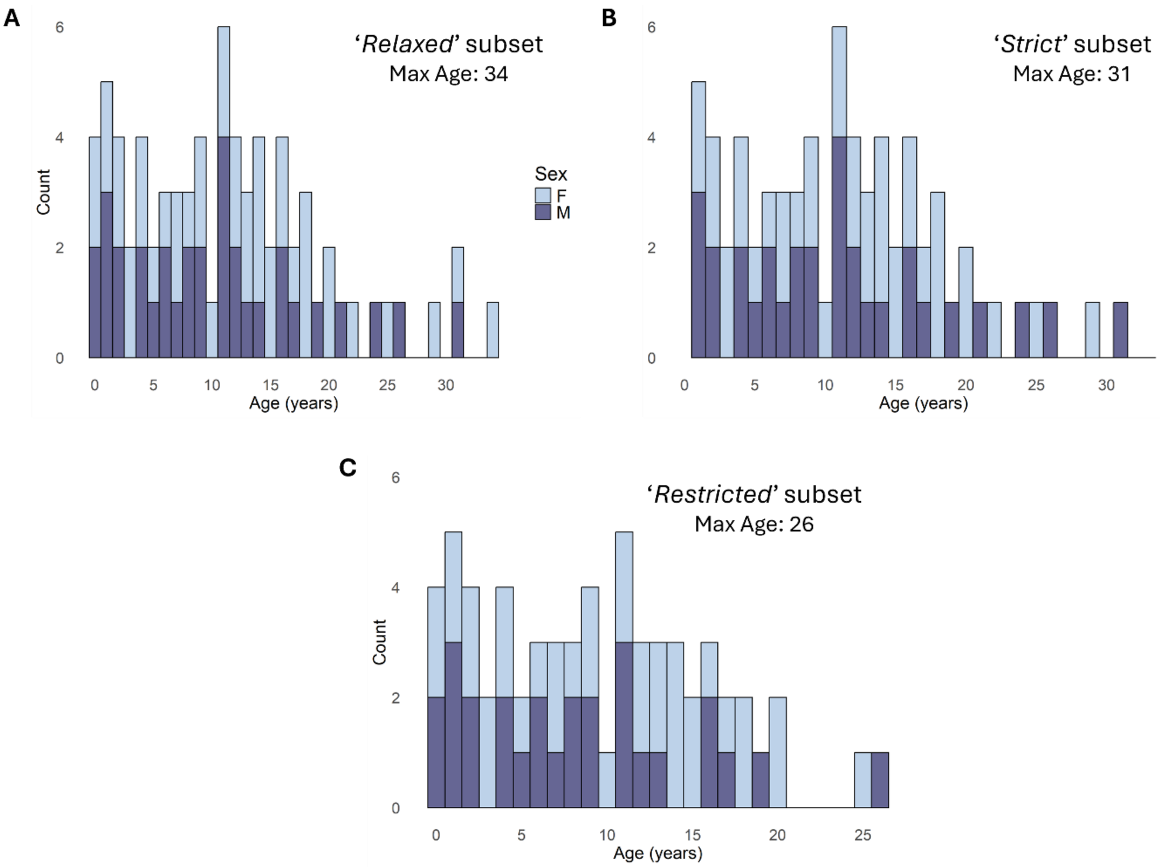


**Figure S1**: Dental age distributions by sex for common dolphins (*Delphinus delphis*) in the **A)** '*relaxed*', **B)** '*strict*', and **C)** '*restricted*' subsets used for epigenetic model development. Max Age = the maximum age in each subset.

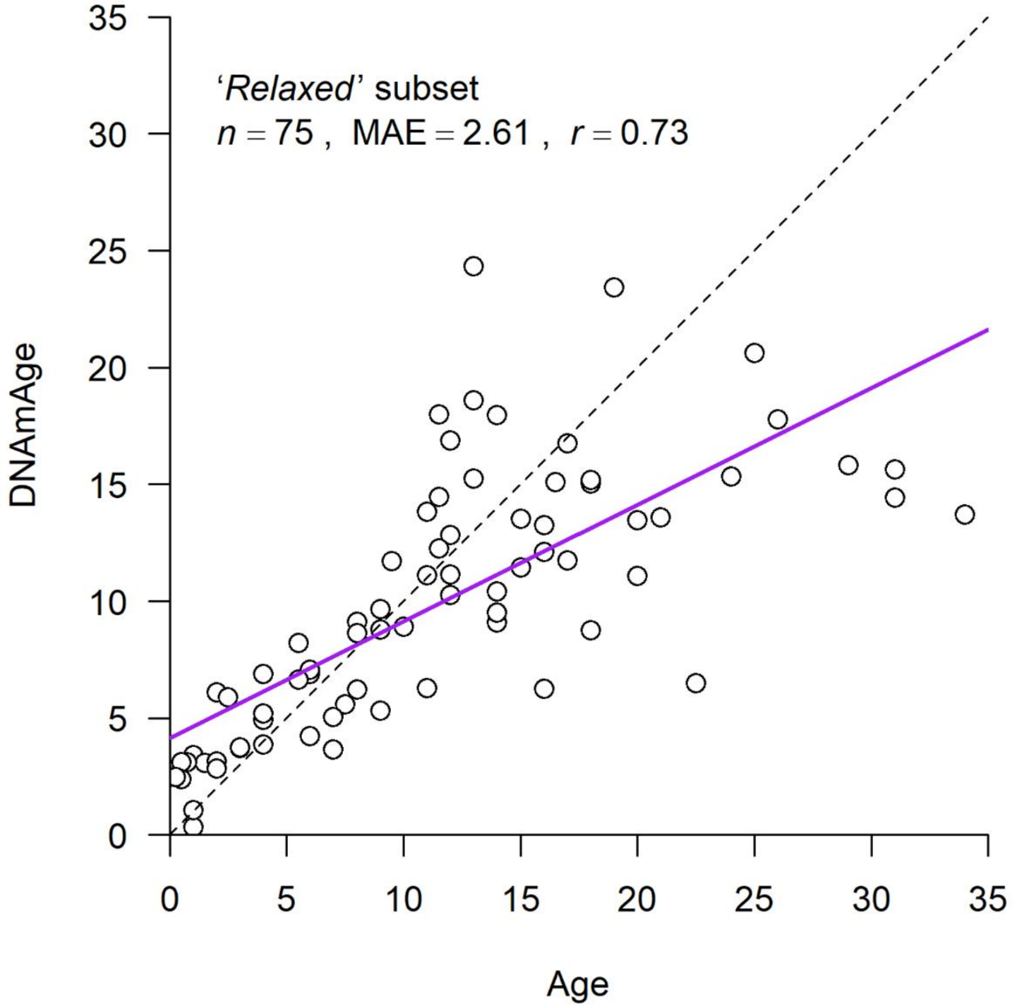


**Figure S2**: Relationship between observed chronological age and DNA methylation-predicted age (DNAmAge) for common dolphin (*Delphinus delphis*) in the ‘*relaxed*’ subset, estimated using a hybrid model. The hybrid approach combined a random forest classifier (RFC) to assign physical maturity status and two elastic net regression (ENR) models: one trained on all individuals and one on physically mature dolphins only. Results are based on 5-fold cross-validation, with RFC and ENR models retrained in each fold to avoid overfitting. Each point represents an individual dolphin; the solid purple line shows the fitted linear regression between observed and predicted ages, and the dashed line marks the 1:1 line of perfect prediction. Model performance metrics (sample size, median absolute error (MAE), and Pearson’s correlation coefficient (*r*) are displayed in the plot title.

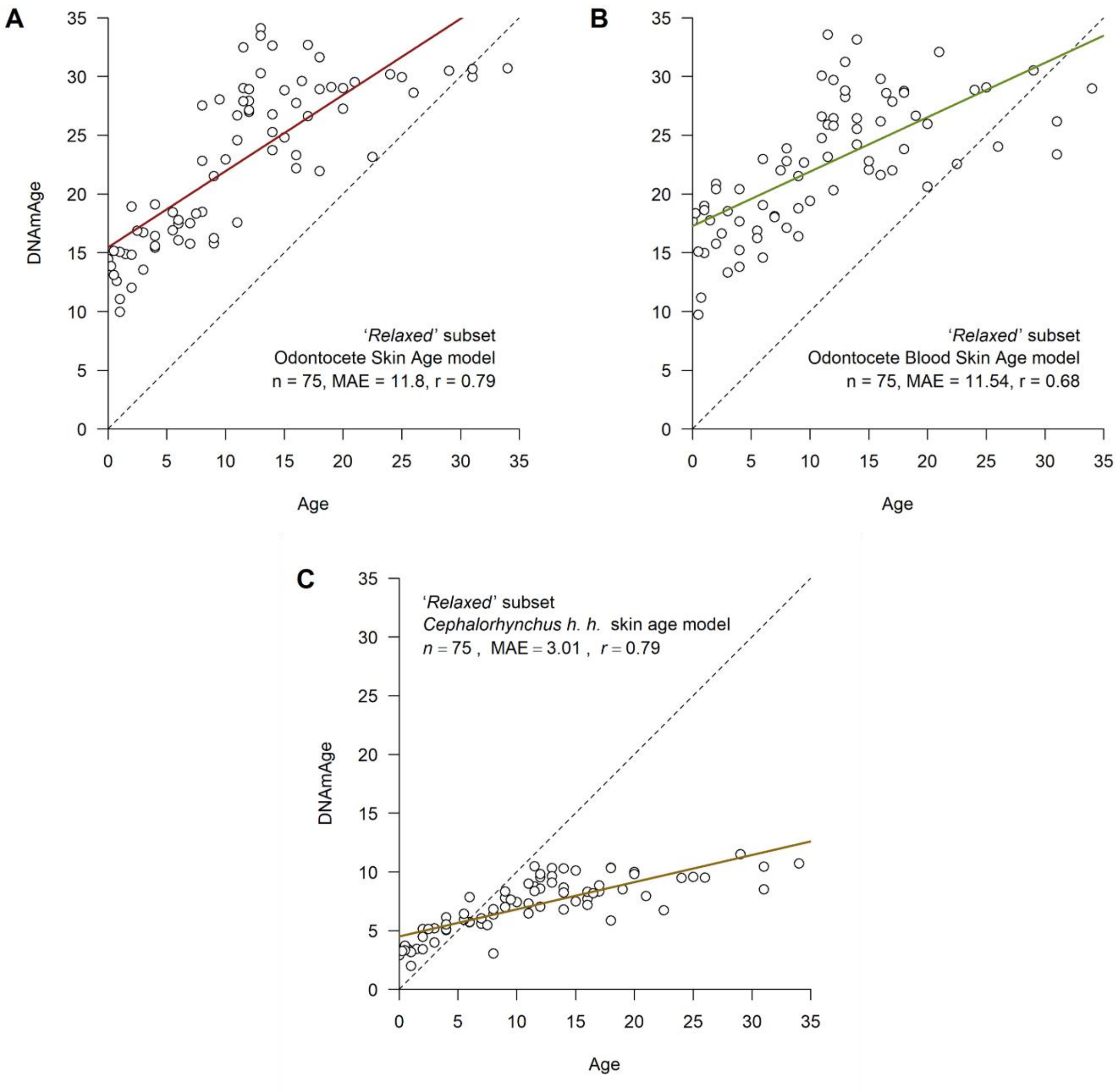


**Figure S3:** Epigenetic ages (DNAmAge) calculated for common dolphin (*Delphinus delphis*) from the '*relaxed*' subset using the **A)** skin clock for multi-species odontocetes (Robeck, Fei, Lu, et al., 2021), **B)** skin and blood clock for multi-species odontocetes (Robeck, Fei, Lu, et al., 2021) and **C)** the *Cetacean Cephalorhynchus hectori hectori* Skin Clock (Hernandez et al., 2023). Regression lines are shown in colour. The dotted diagonal indicates a perfect correlation with y = x. Individual animals are represented by green dots. Pearson correlation (r) and median absolute error are given for each model.
